## Supplementary figures and images for "Tracing the evolution and genomic dynamics of mating-type loci in *Cryptococcus* pathogens and closely related species"

### S1 Fig

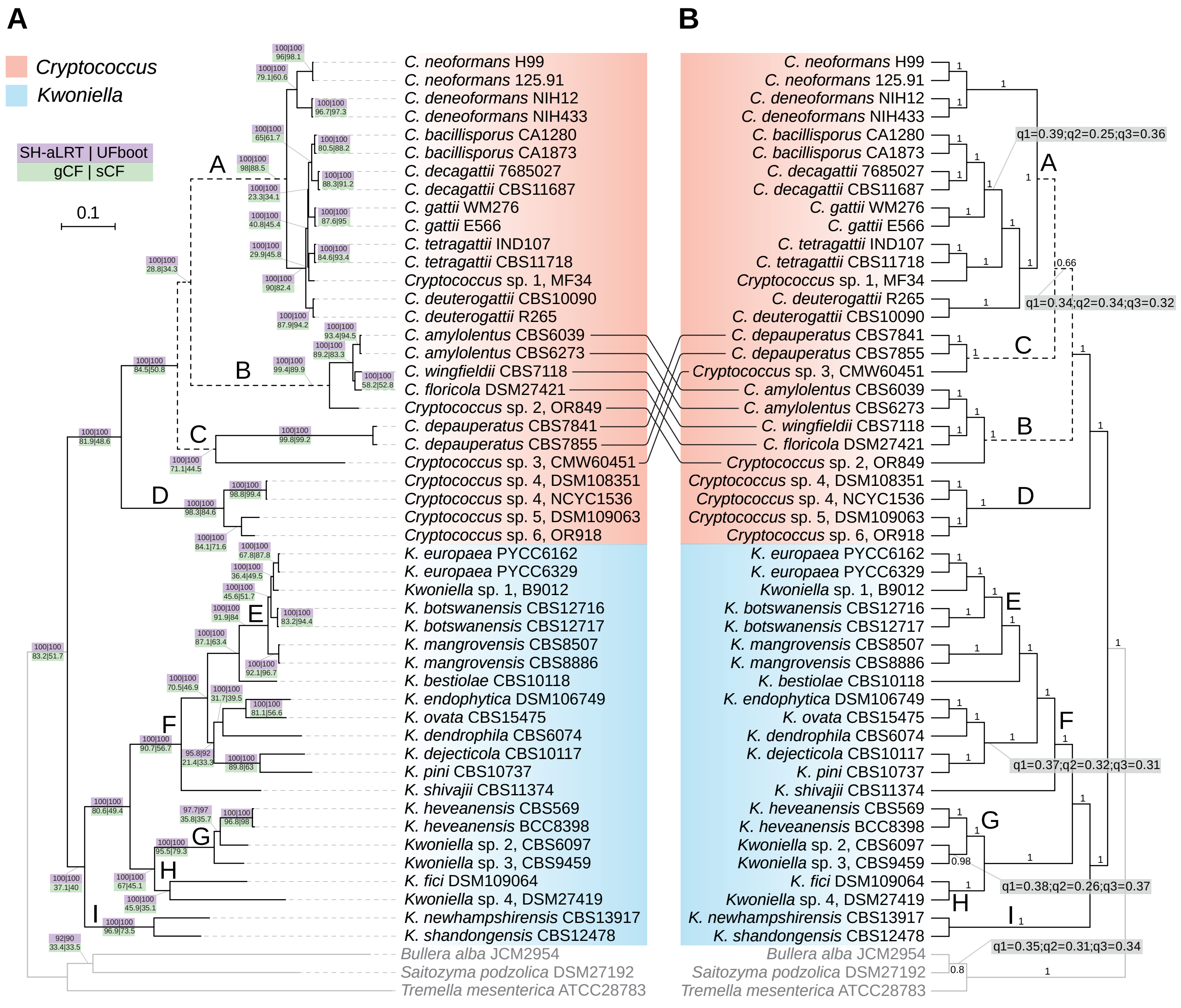

### S2 Fig

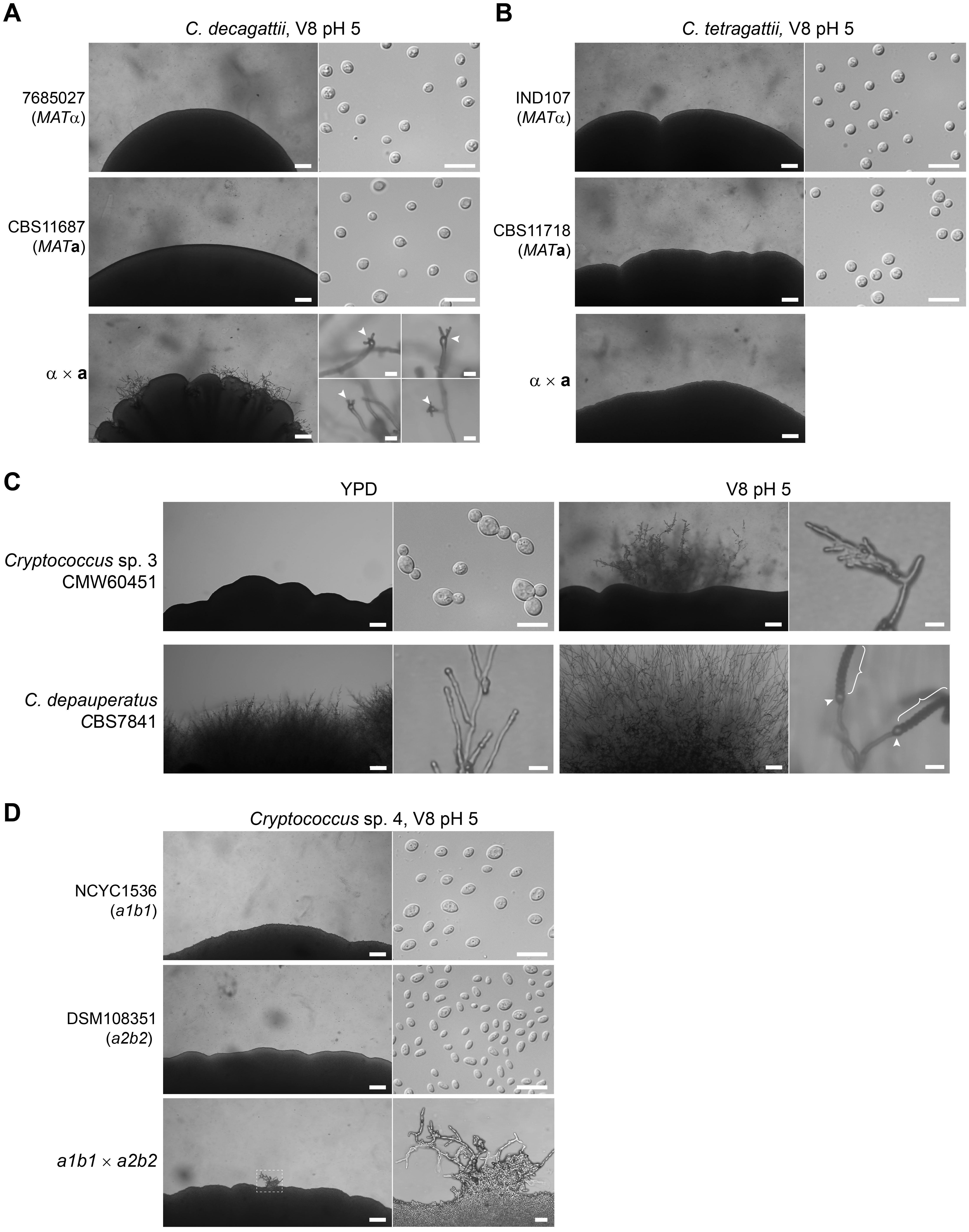

### S3 Fig

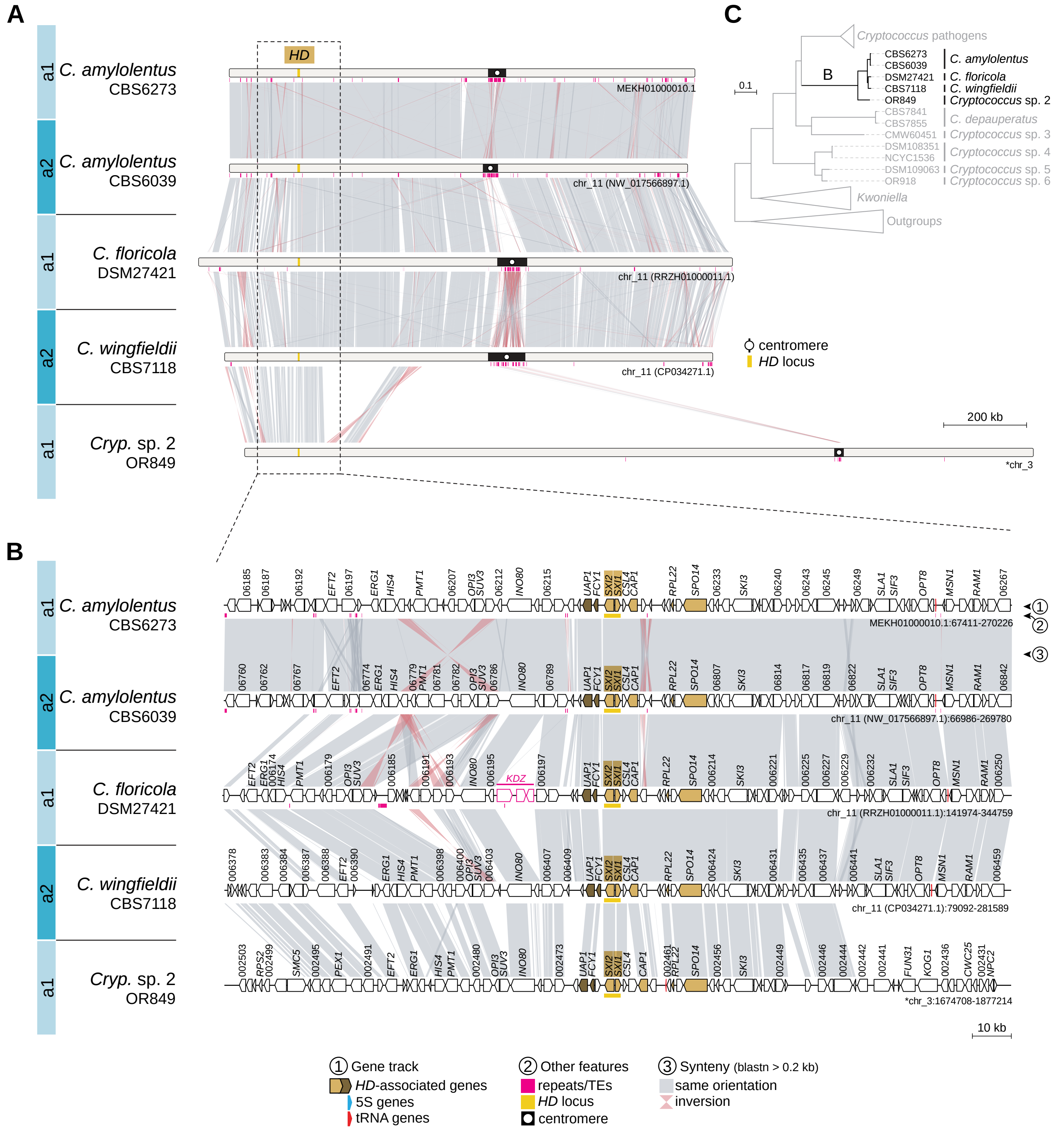

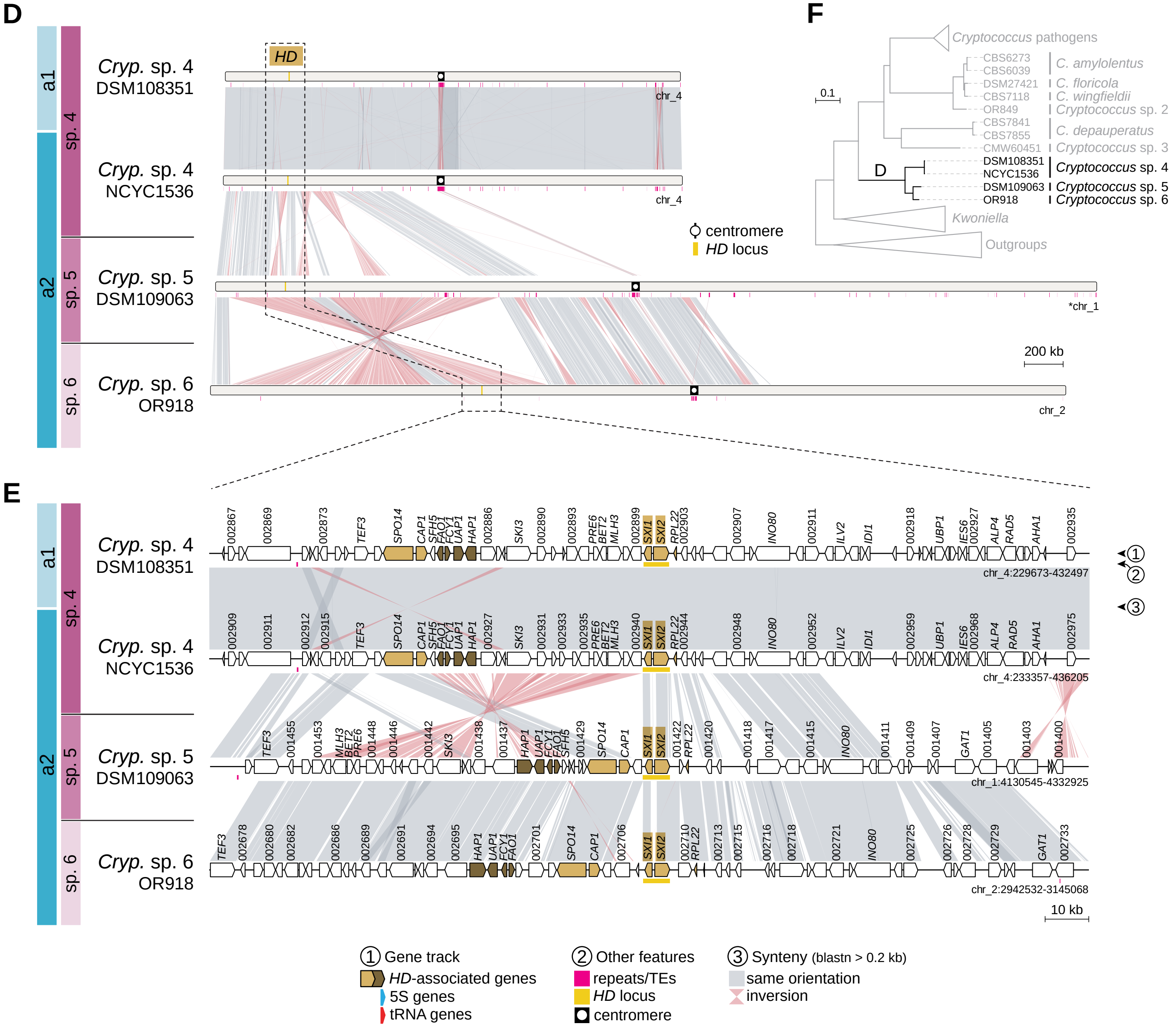

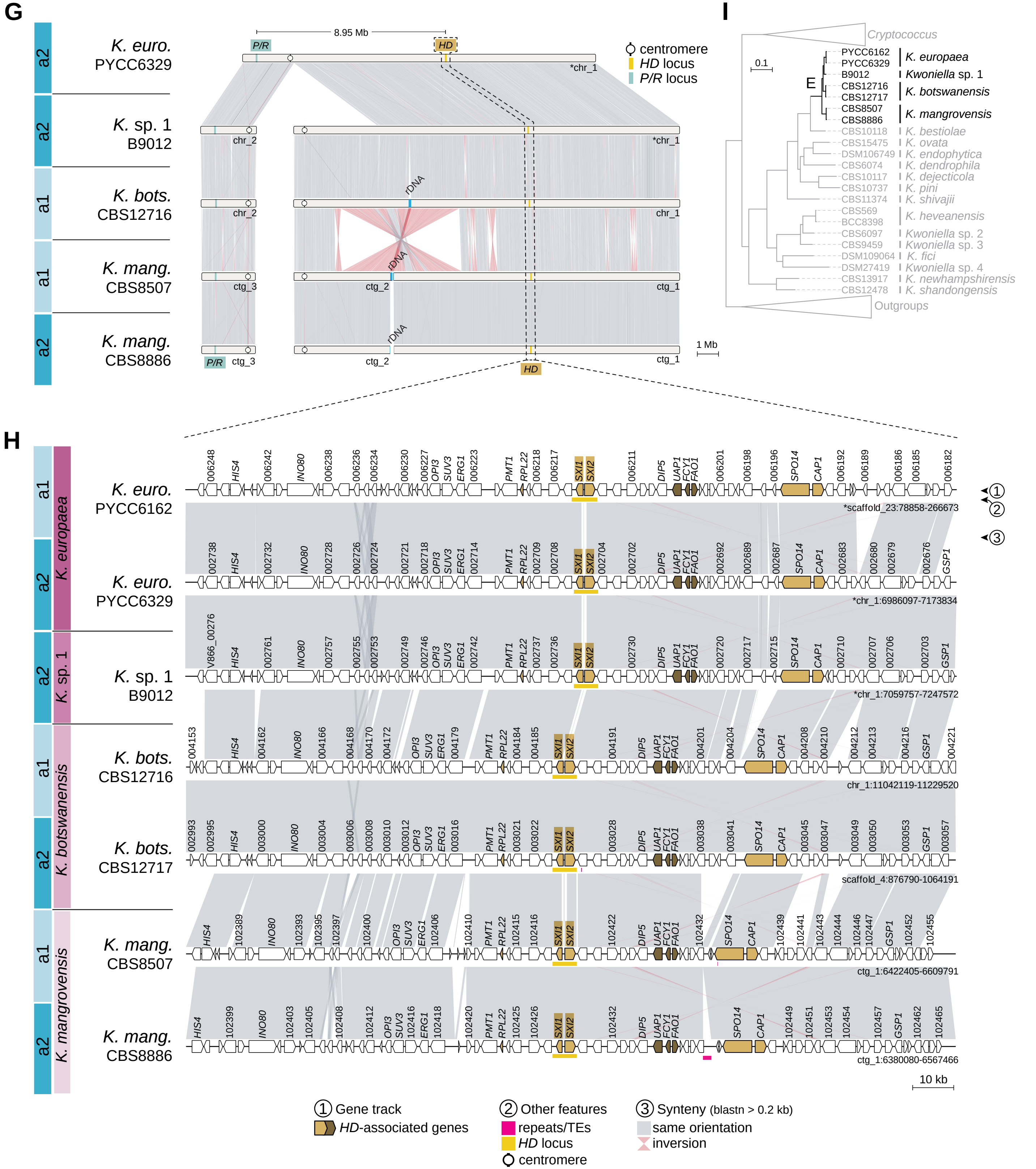

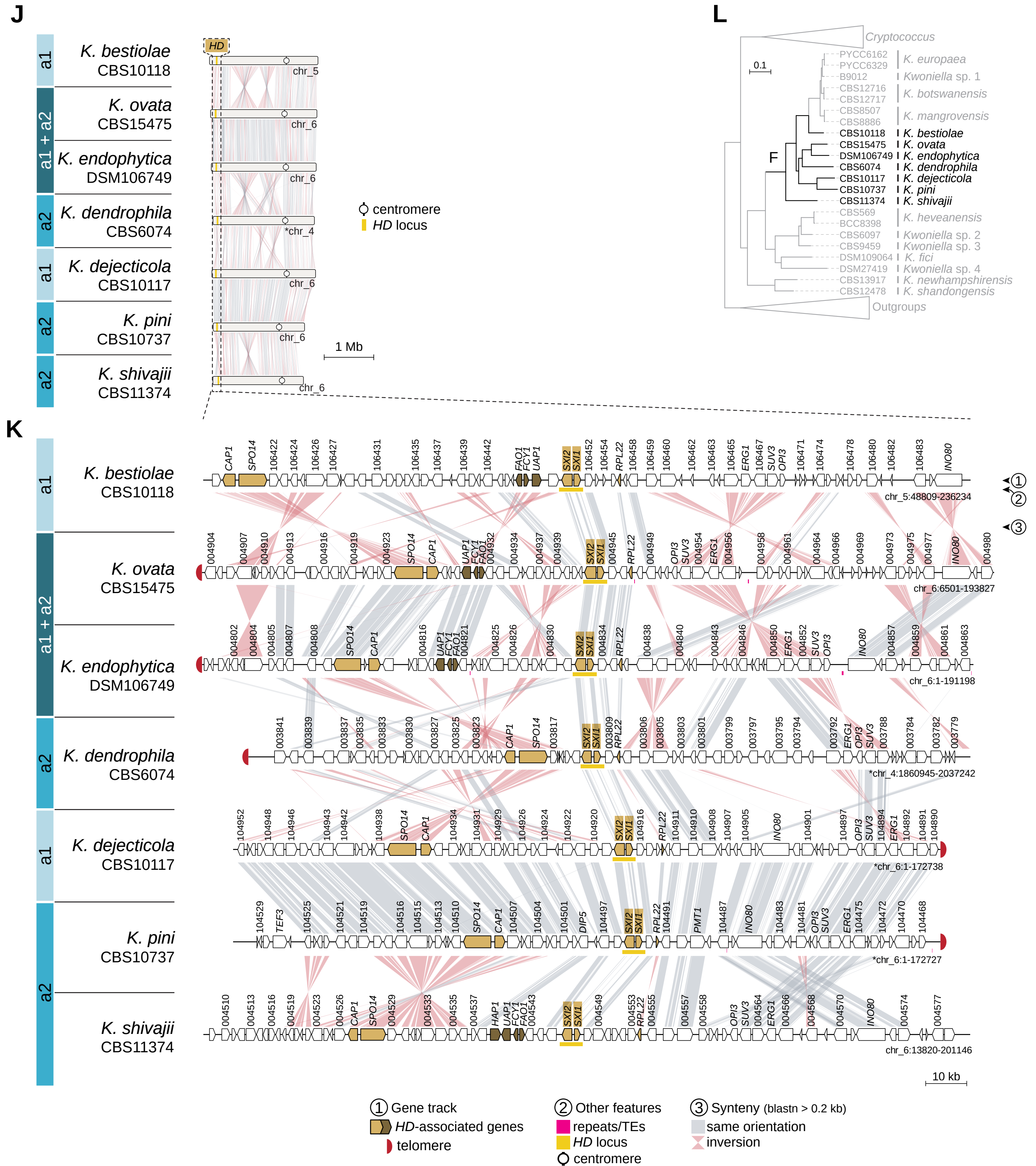

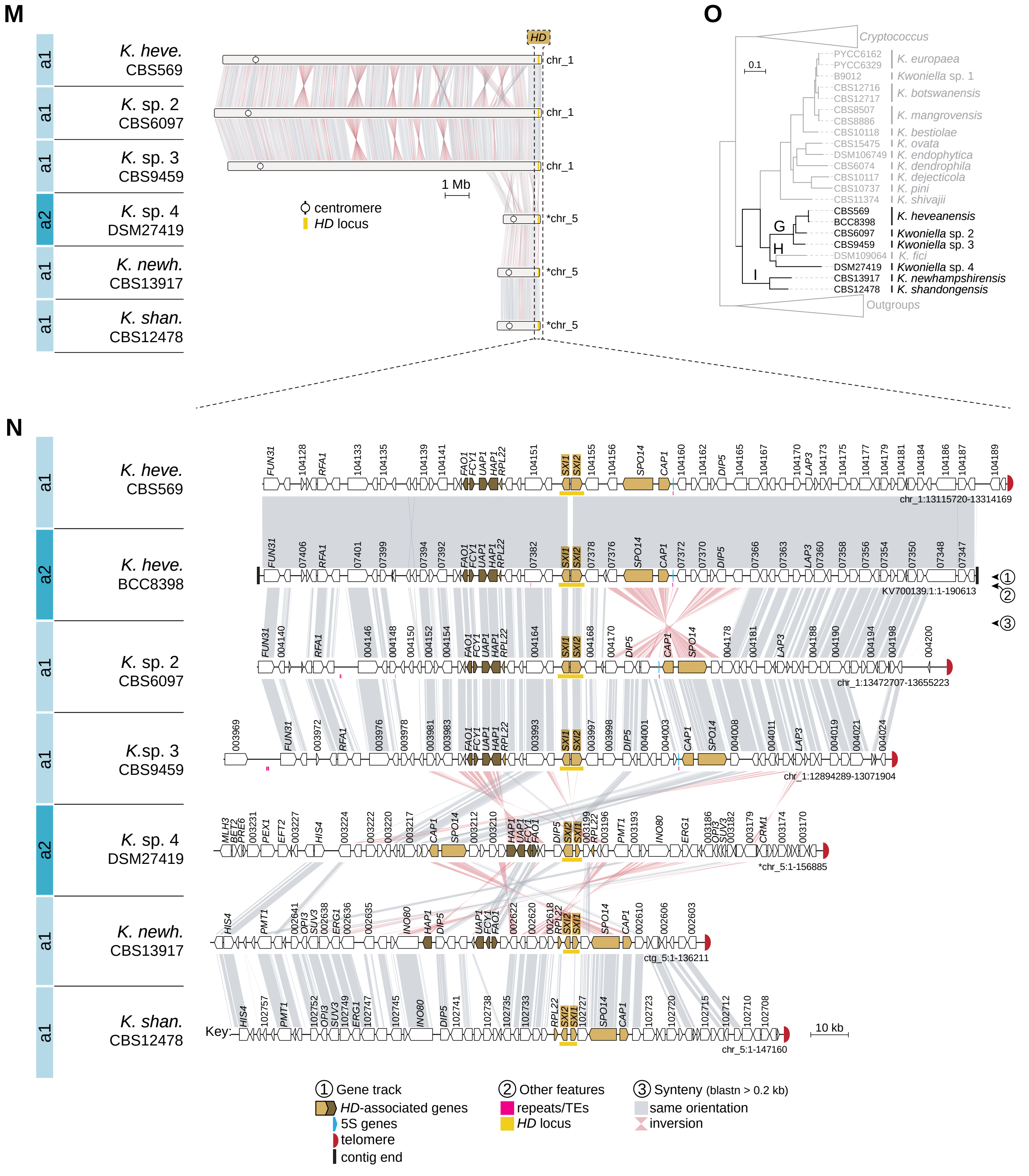

### S4 Fig

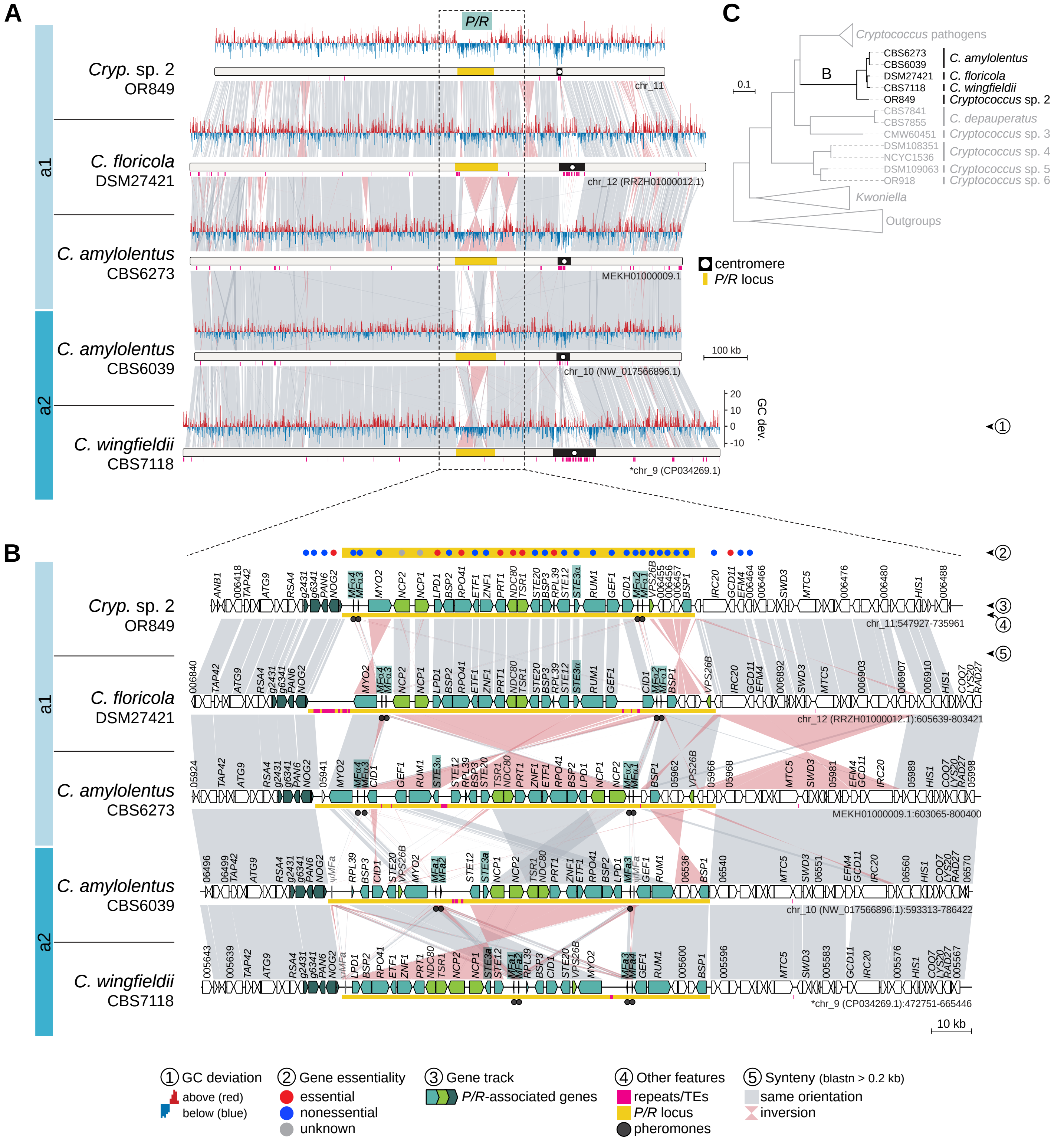

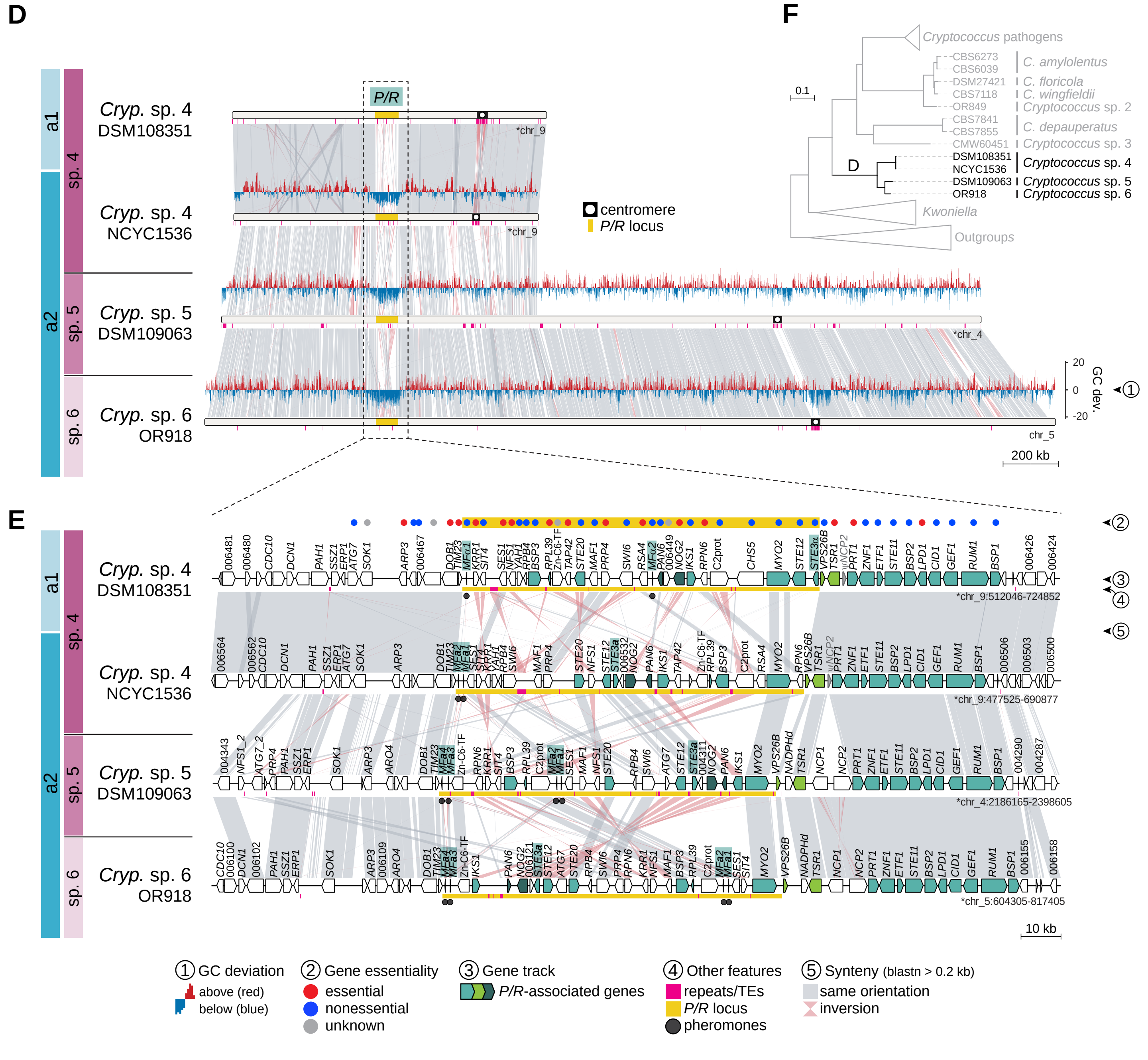

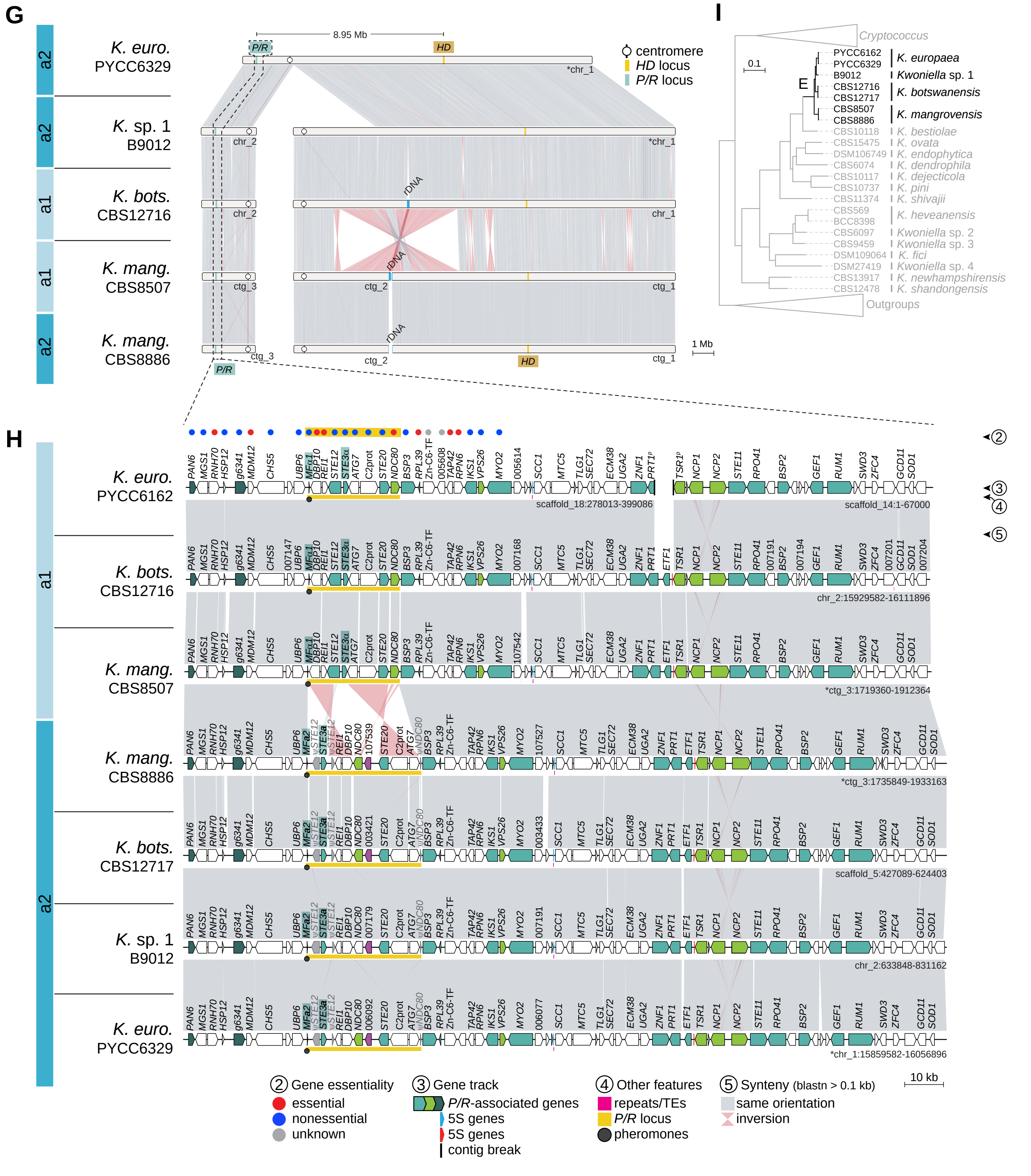

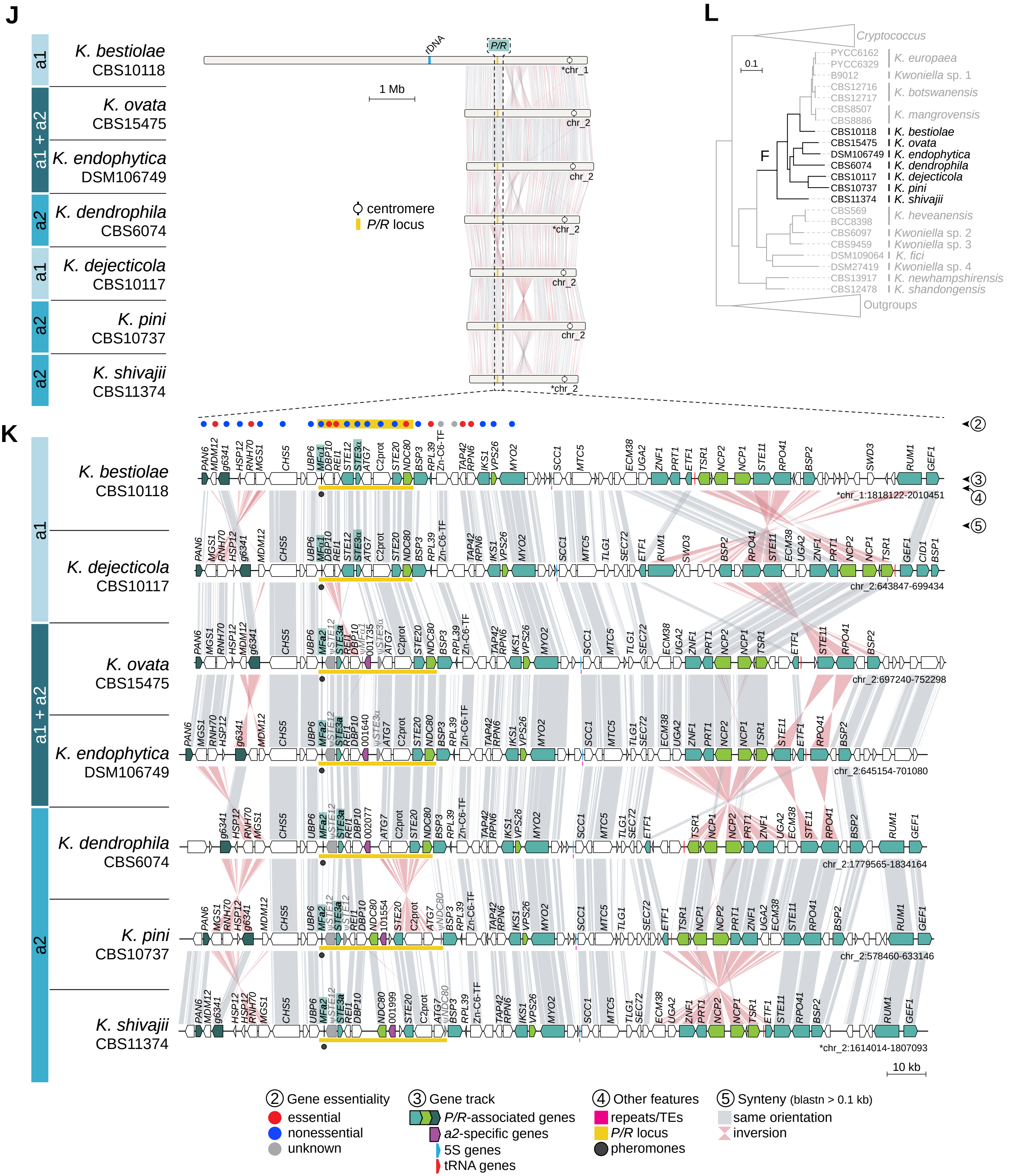

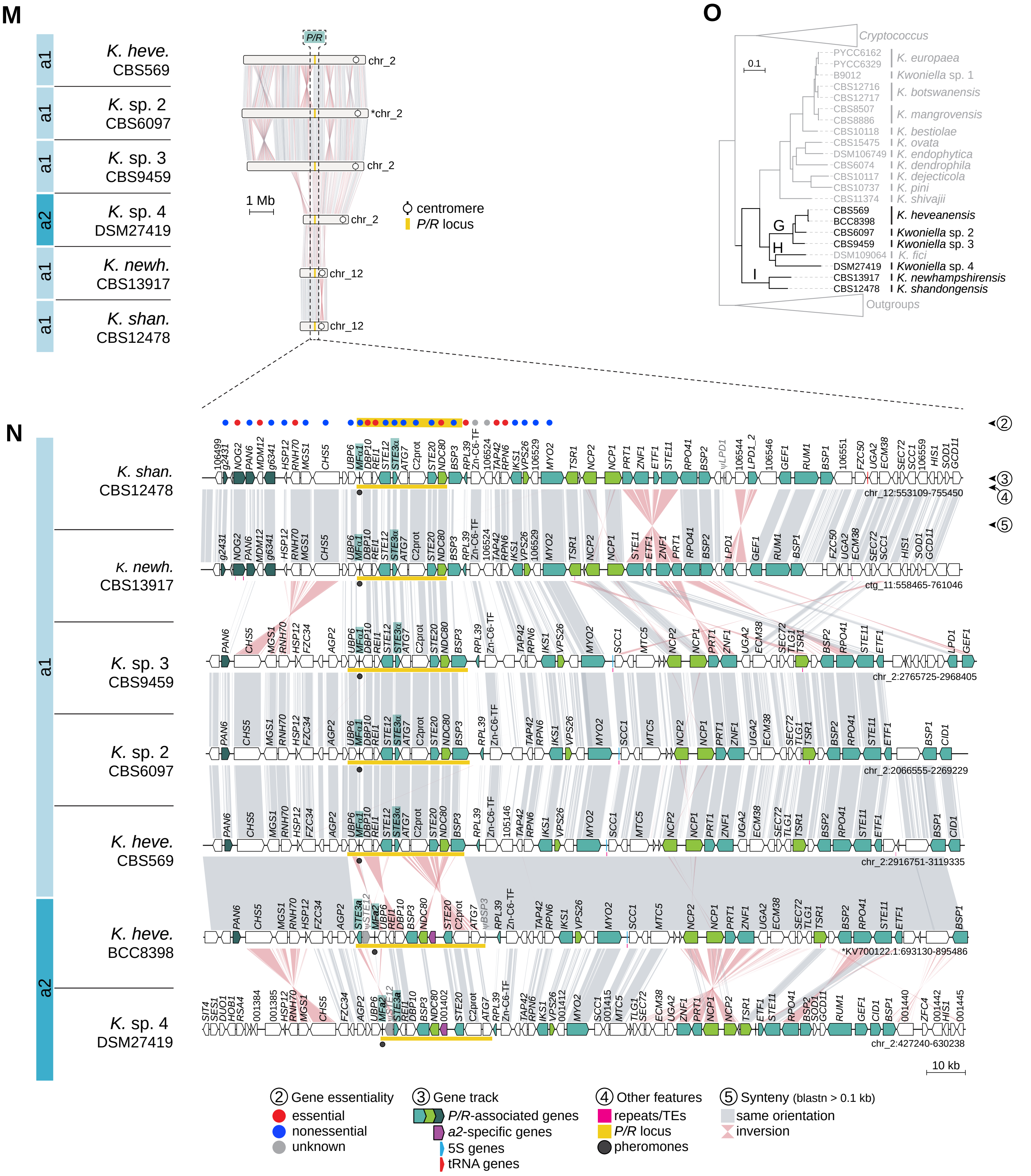

### S5 Fig

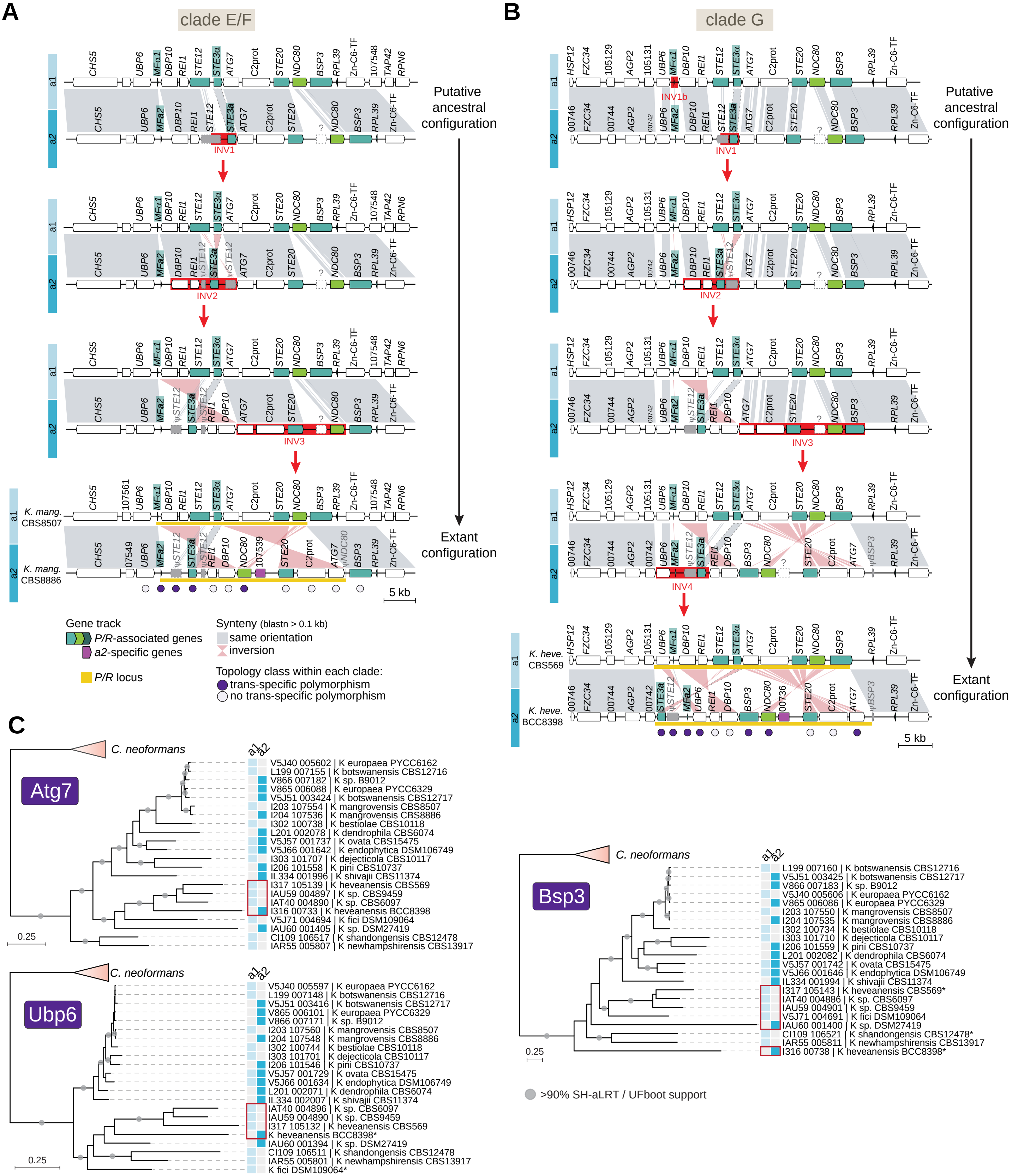

### S6 Fig

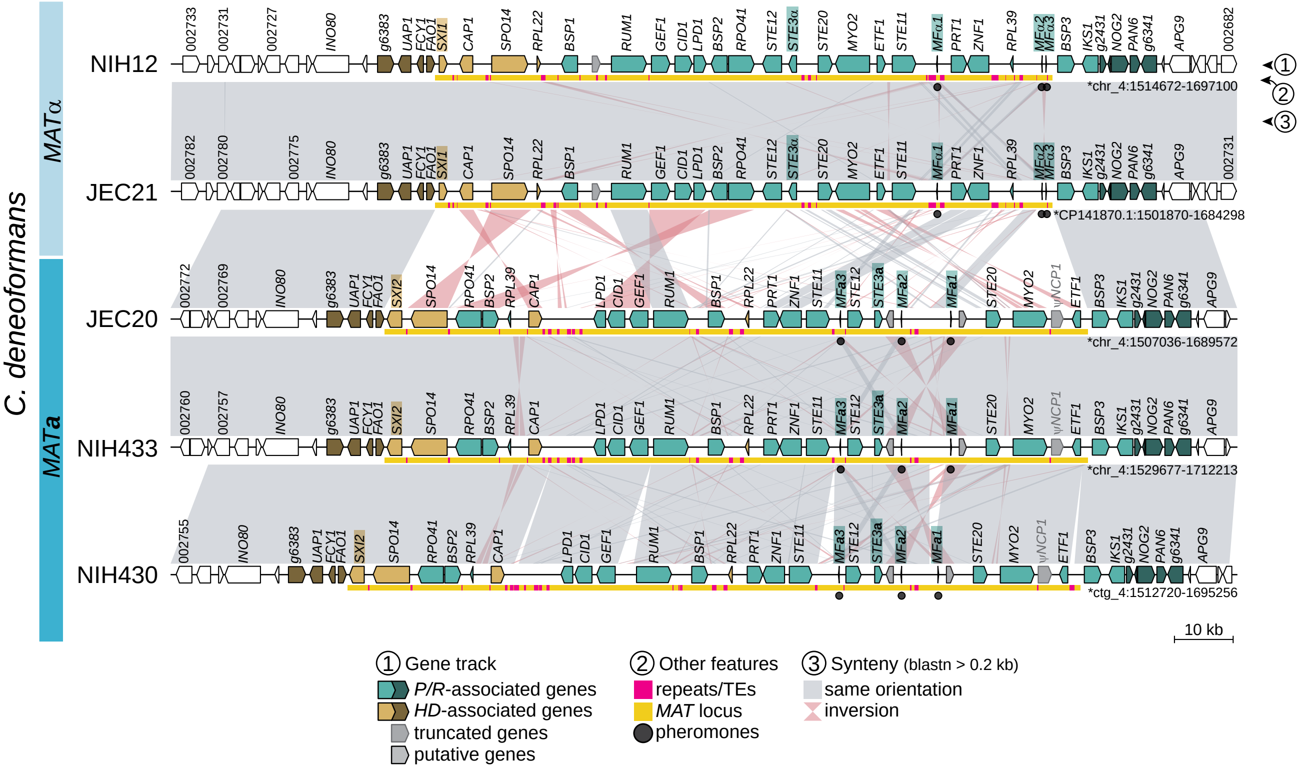

### S7 Fig

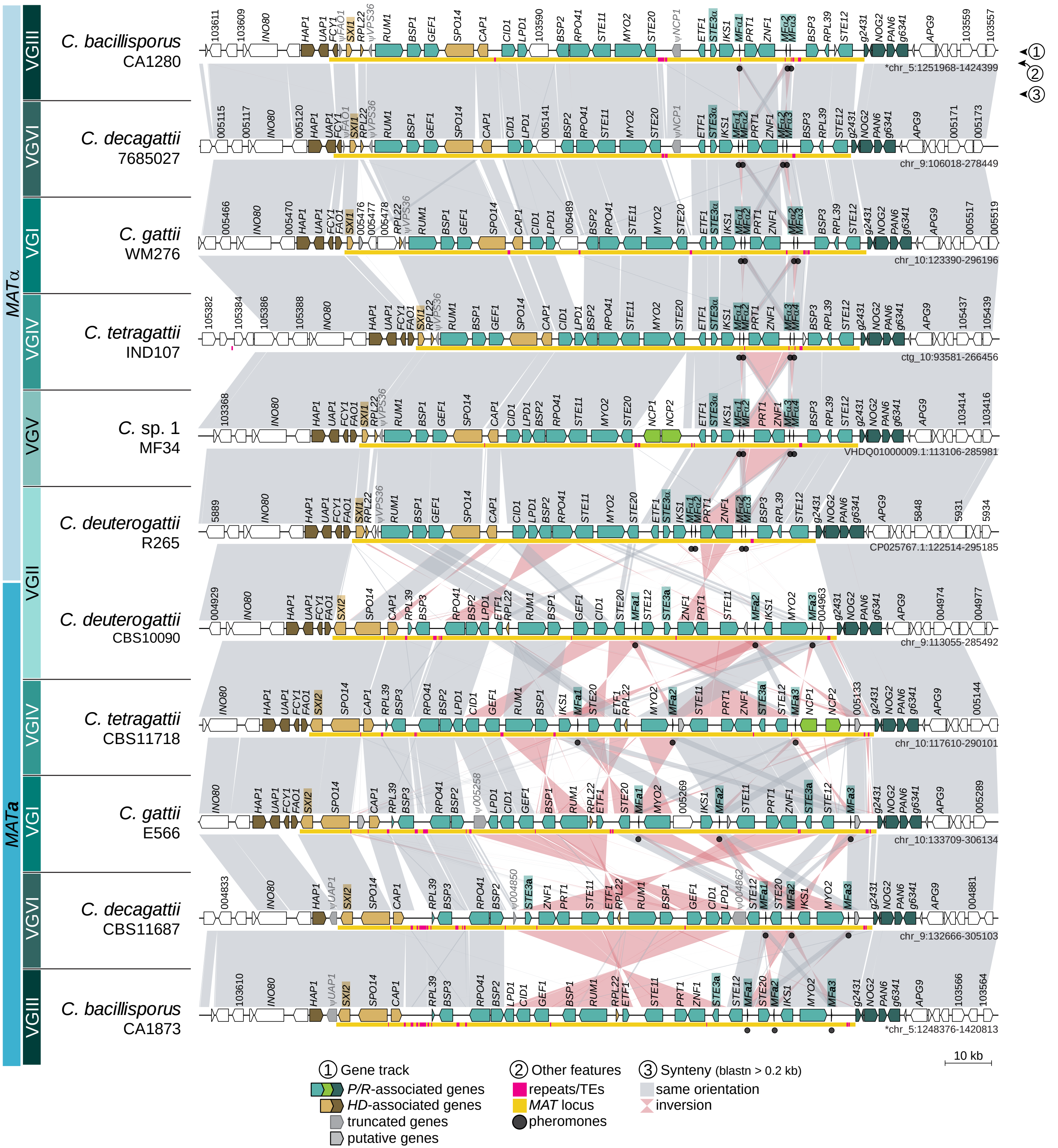

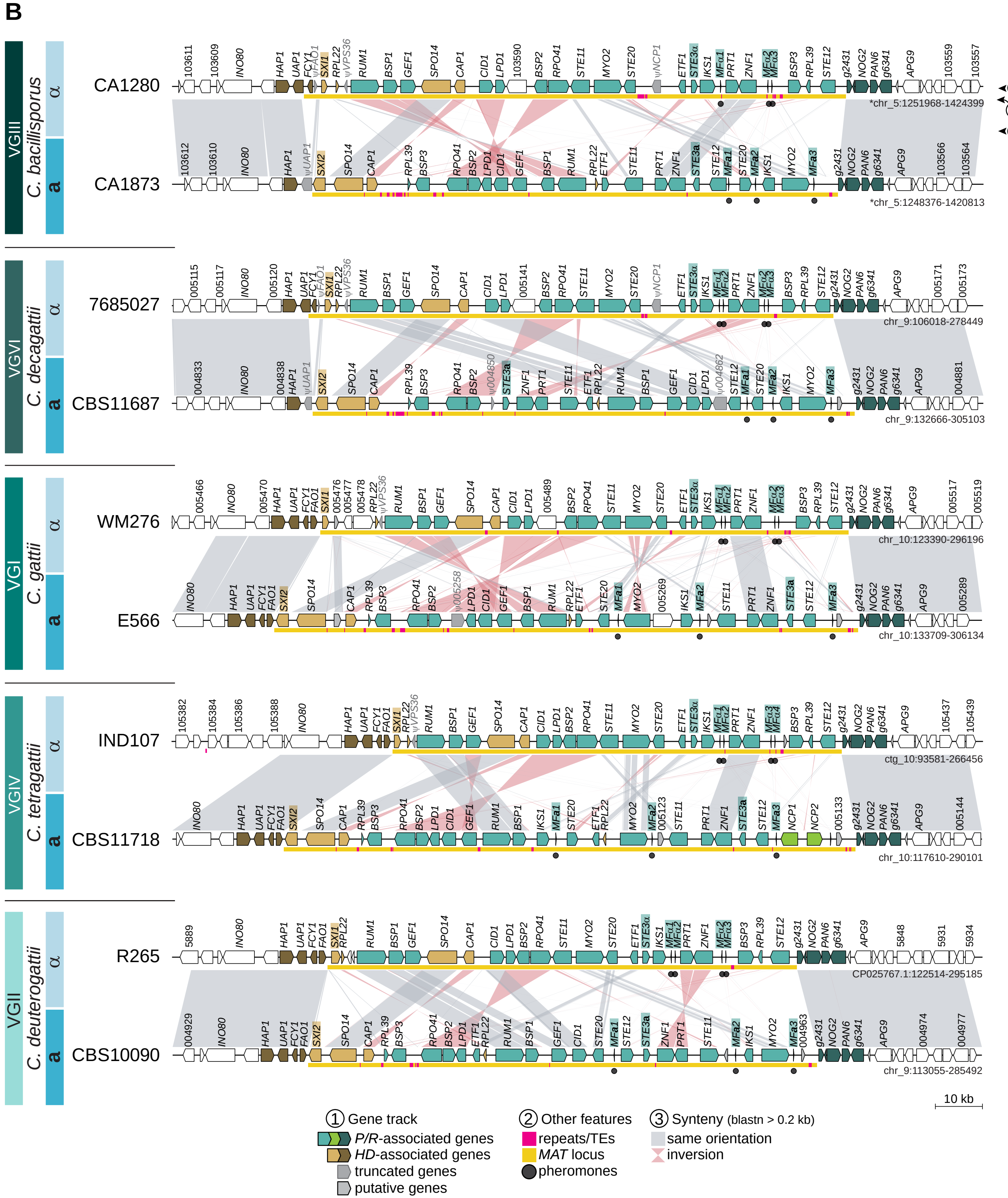

### S8 Fig

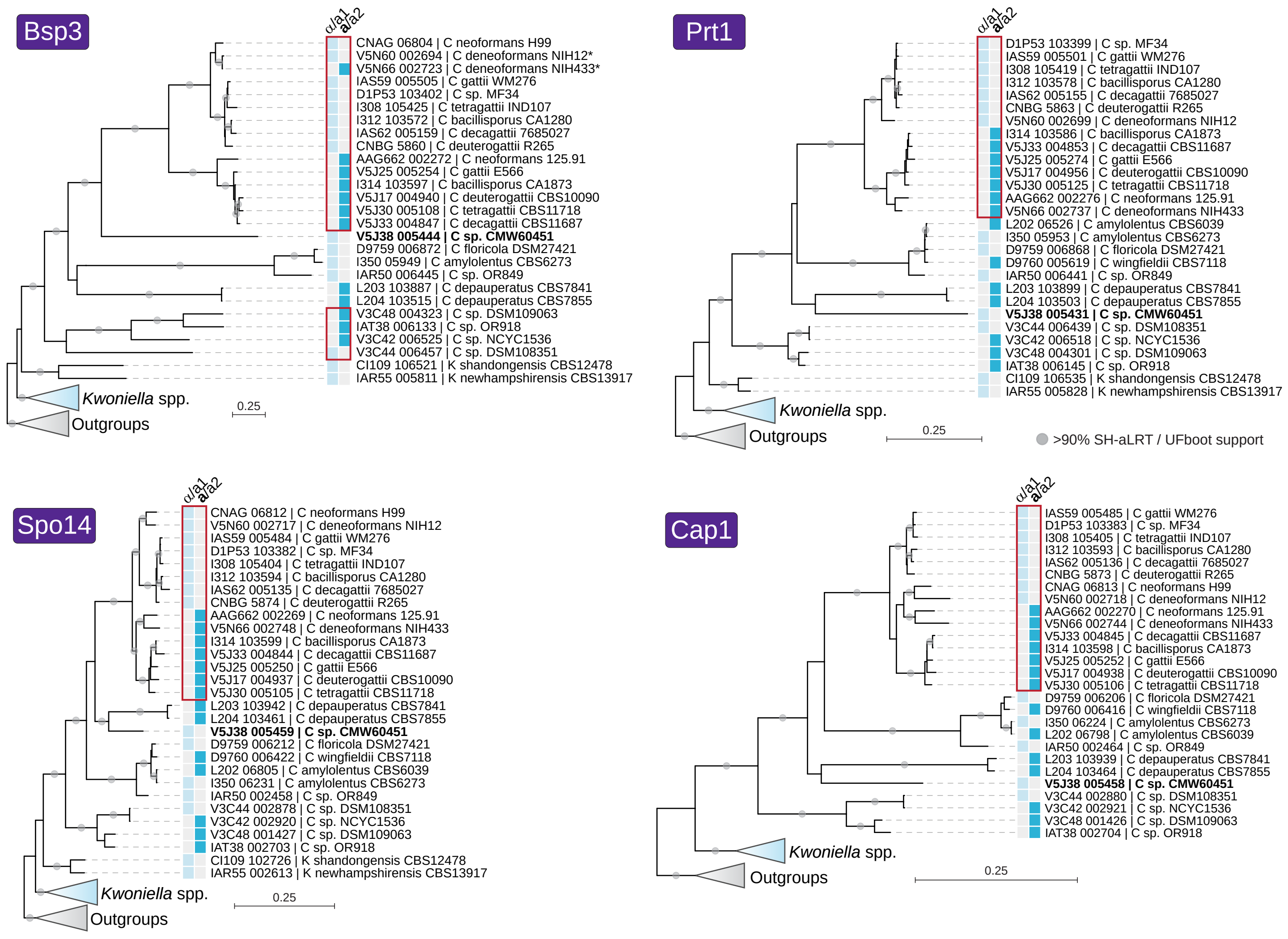

### S9 Fig

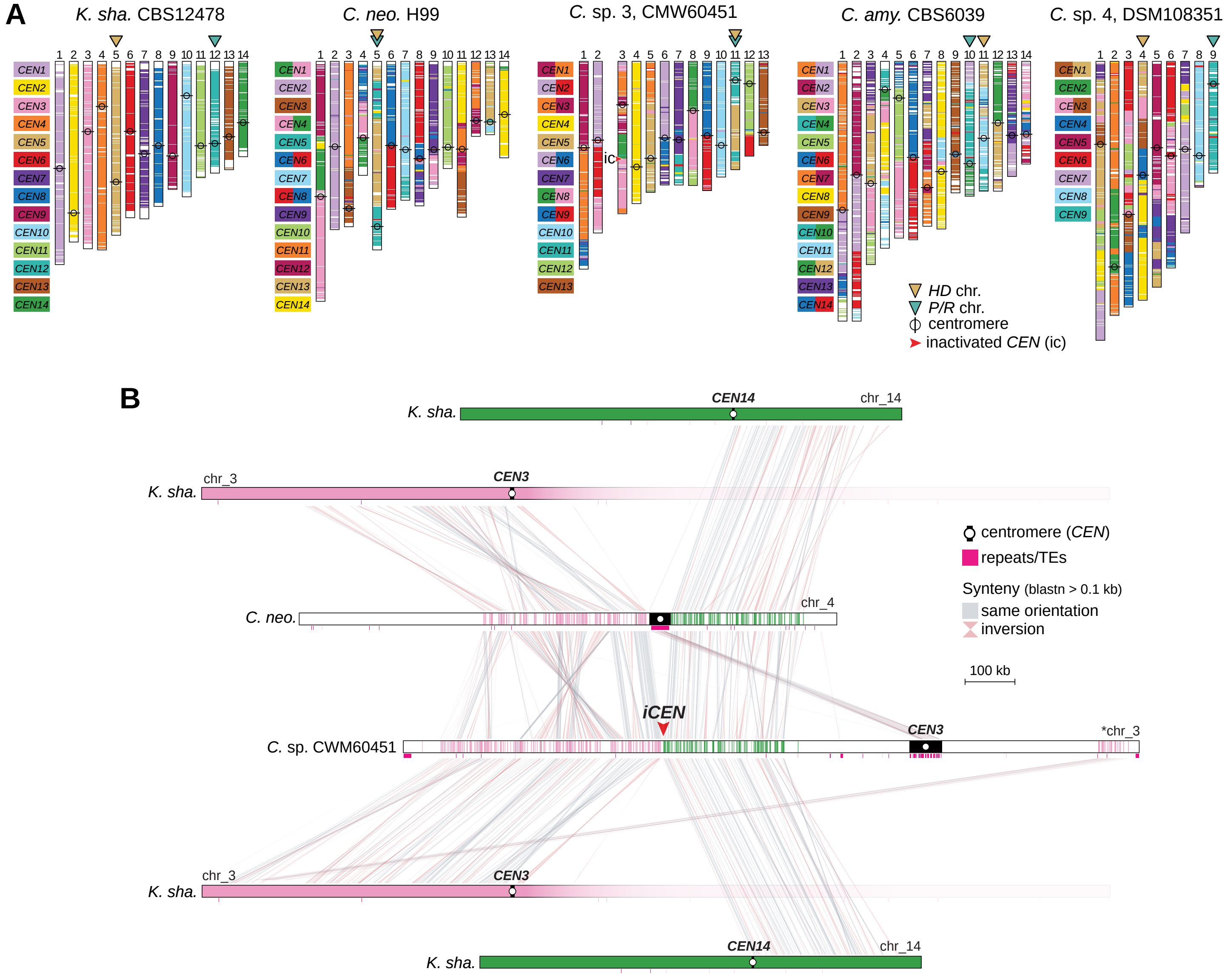

### S10 Fig

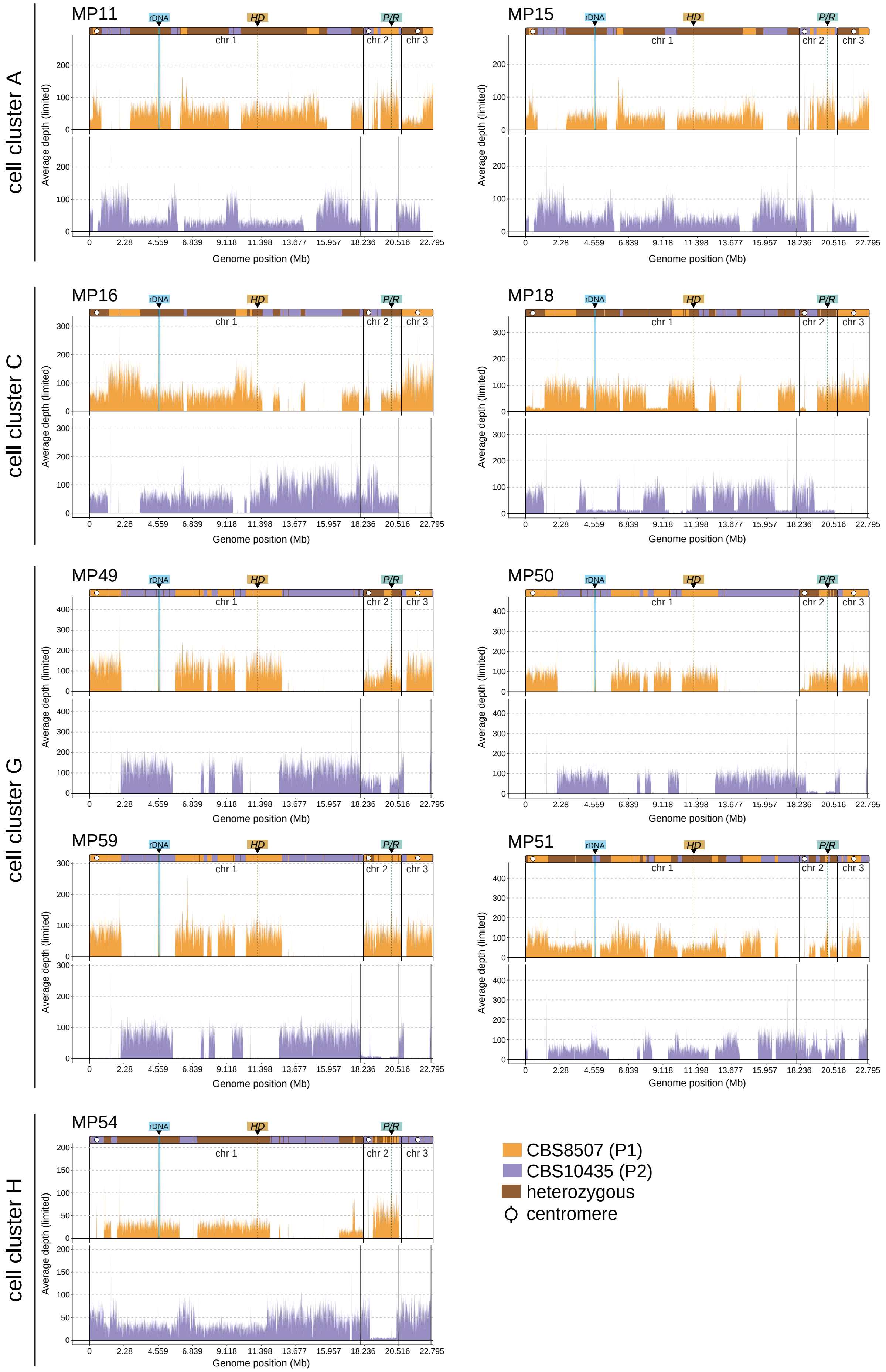

### S11 Fig

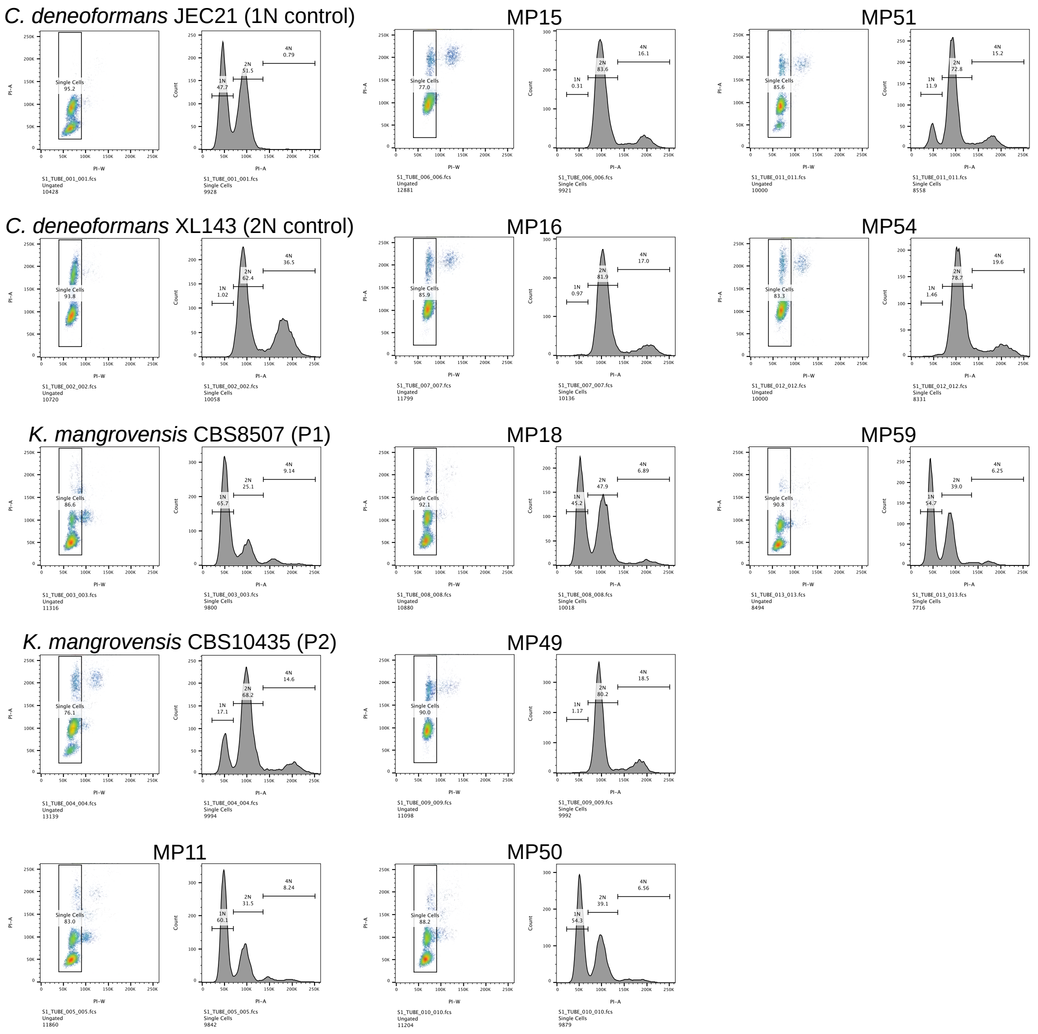

### S12 Fig

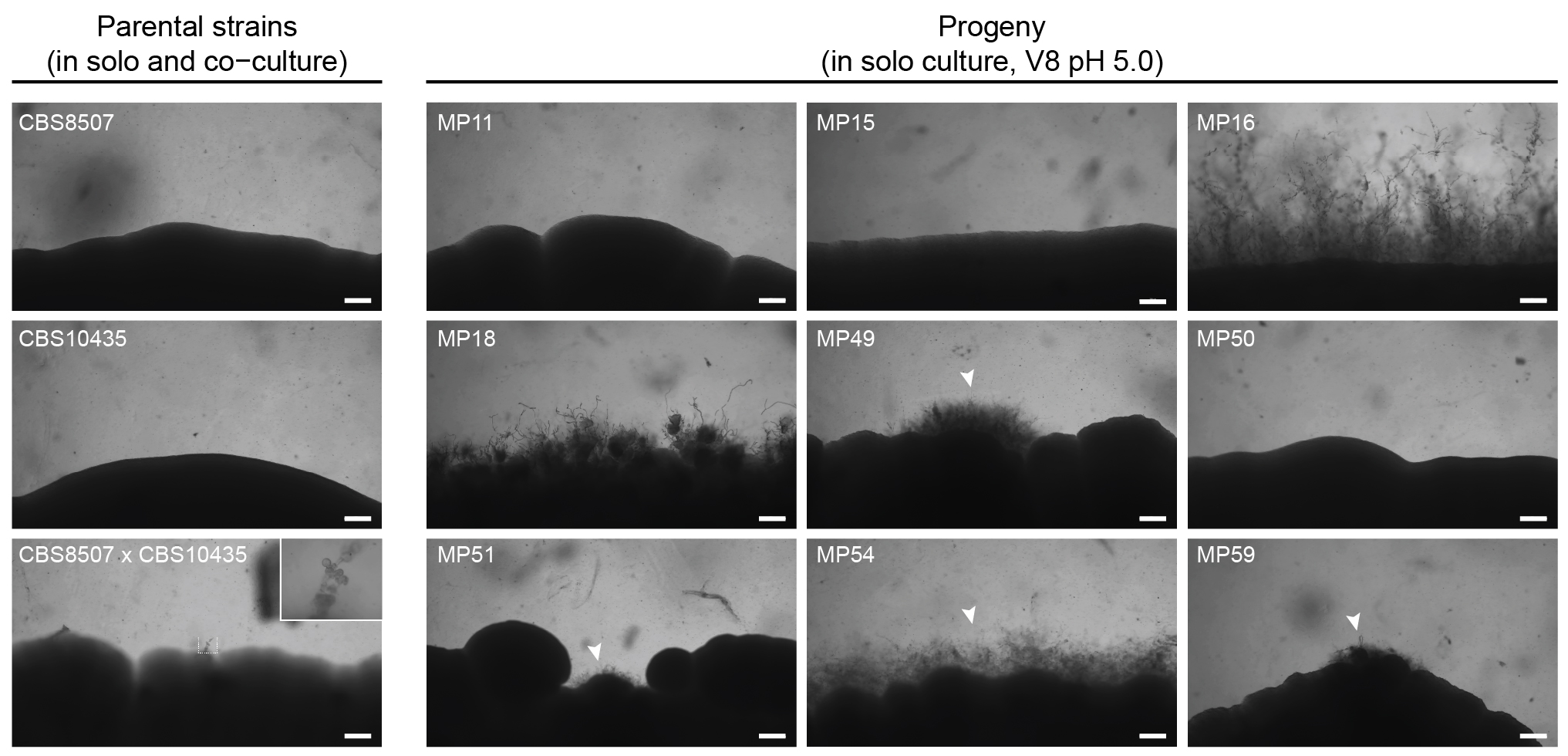
