## Supplementary material for "Tracing the evolution and genomic dynamics of mating-type loci in *Cryptococcus* pathogens and closely related species": S1 Text

### Overview and rationale

We evaluated whether the lower GC content of the pheromone/receptor (*P/R*) locus in clades B and D mainly reflects a neutral consequence of reduced recombination, specifically reduced GC-biased gene conversion (gBGC), or whether weaker selection on synonymous codon usage also contributes. gBGC is a recombination-associated fixation bias that tends to favor G/C alleles and elevate GC content in regions where recombination is frequent [1, 2]. If recombination is suppressed, this GC-increasing effect weakens, and sequence composition drifts toward AT. In parallel, codon usage can be influenced by selection for translational efficiency and accuracy [3, 4]; if this selection is relaxed, codon usage can move closer to background nucleotide composition.

To disentangle these effects, we performed four complementary analyses that compare GC content and codon usage patterns in the *P/R* region versus matched regions outside *P/R*, allowing us to separate neutral shifts driven by recombination-associated gBGC from those potentially shaped by selection. Specifically, we conducted: (1) a neutral comparison of GC content in non-coding regions (intergenic plus intronic sequence) within the P/R locus versus length-matched, *P/R*-excluded windows on the same chromosome; (2) an analysis of third-position nucleotide composition in coding regions (AT3 vs. GC3), which reflects both gBGC and synonymous codon selection; (3) codon-bias index comparisons between *P/R* and background (BG) genes, including frequency of optimal codons (Fop), observed effective number of codons (Nc; [5]), the expected Nc at the observed GC3 (ENC), and ΔENC = Nc − ENC; and (4) codon adaptation index (CAI) analyses that explicitly control for GC3 and CDS length, to test whether codon usage deviates from compositional expectations.

The logic is that non-coding DNA provides a neutral baseline for gBGC signals, whereas coding regions may also reflect selection on synonymous codons; conditioning CAI (and Fop) on GC3 and length helps distinguish composition from selection on translation. Unless stated otherwise, comparisons are between genes fully contained within the *P/R* window for each strain and same-chromosome genes outside the *P/R* region. Raw metrics and column names (referenced in parentheses and italics) are given in **S4 Appendix**.

#### (1) Neutral non-coding GC (intergenic + introns)

We compared non-coding GC within the *P/R* window (*gc_intergenic*; intergenic plus intronic sequence) to a same-chromosome, length-matched null built from n = 10,000 *P/R*-excluded windows (*null_mean_intergenic*). Non-coding sequence is not shaped by selection on synonymous codons and is therefore a standard baseline for detecting composition shifts attributable to gBGC rather than selection on translation [2, 6]. For each strain, we first placed as many non-overlapping windows as possible with ≥1 kb spacing. Because chromosomes cannot accommodate 10,000 disjoint windows of ~85–100 kb, we then sampled additional *P/R*-excluded starts with replacement at base-pair resolution to reach n = 10,000, allowing overlap among null windows. We report ΔGC (*delta_gc*) = *gc_intergenic* − *null_mean_intergenic*, a standardized z score (*z_intergenic*) from the null distribution, and a one-sided empirical permutation p-value (*perm_p_intergenic*) for P/R ≤ null.

This analysis showed a strong neutral signature in clade D across all species (ΔGC ≈ −0.056 to −0.083; permutation test, *P* = 1e−4; z ≤ −5). In clade B, depletion was smaller and often not significant (e.g., CBS6039 ΔGC ≈ −0.0038, perm. *P* = 0.27), with one exception (*C. wingfieldii* CBS7118: ΔGC ≈ −0.0177, perm. *P* = 1e−4; z = −2.16). These patterns are consistent with reduced gBGC in a recombination-suppressed *P/R* region, particularly in clade D.

#### (2) Third-position composition inside coding regions (AT3 vs. GC3)

We next asked whether coding sequences show the expected shift under weaker gBGC by tallying third positions that end in A/T vs. G/C inside *P/R* (*AT3_PR/GC3_PR*) and in same-chromosome background (*AT3_BG/GC3_BG*), and evaluating 2×2 AT3/GC3 contingency tables (*p_endings*; $\chi^2$ with Fisher's exact test fallback). Third codon positions are largely synonymous and thus sensitive to both background nucleotide composition and gBGC, allowing detection of AT3↔GC3 shifts without altering protein sequence [7]. We also compared per-gene GC3 summaries (e.g., *median_GC3*).

Both clades exhibited extremely significant shifts toward AT3 inside the *P/R* locus (clade D *p_endings* $\approx 0$ for all species; clade B *p_endings* $\ll$ 1e−70), with a larger GC3 drop in clade D (e.g., DSM108351 *GC3_PR* = 7,514 vs. *GC3_BG* = 157,693) than in clade B (e.g., CBS6039 *GC3_PR* = 9,388 vs. *GC3_BG* = 136,267). Per-gene summaries agreed (e.g., DSM108351 *median_GC3* = 0.499 inside vs. 0.749 in background; CBS6039 0.556 vs. 0.626). Thus, coding sequences inside *P/R* are AT3-biased relative to background, consistent with weaker gBGC and aligning with the neutral non-coding result.

#### (3) Codon-bias indices (selection-sensitive)

To evaluate selection on synonymous codon usage beyond composition, we computed Fop, Nc, ENC, and ΔENC = Nc − ENC for each gene. These indices are standard tools for quantifying translational selection [5, 8]. Fop measures enrichment of "optimal" codons that match abundant tRNAs; Nc summarizes codon diversity (values near 20 indicate strong bias; values near 61 indicate little/no bias). ΔENC compares observed Nc to the expectation at the observed GC3, helping separate true codon preference/optimization from composition (i.e. patterns explained just by GC content) [9]. We contrasted *P/R* vs. background by median differences with Mann–Whitney and permutation tests (*Fop_med_diff/Fop_perm_p*; *Nc_med_diff/Nc_perm_p*; *DeltaENC_med_diff/DeltaENC_perm_p*), emphasizing medians because Nc can be noisy for short ORFs. Because Nc ranges from 20 (strong bias) to 61 (no bias; [5]) and can be unstable for short genes, we applied conservative adjustments to ensure that comparisons reflect real biological trends rather than technical noise, specifically keeping Nc within this natural range (details in **Materials and Methods**); ΔENC was then computed from this Nc and ENC.

In clade D, *P/R* genes used fewer optimal codons (negative *Fop_med_diff*, all permutation tests: $P \leq$ 1e−4), but Nc medians frequently tied at 61 in both *P/R* and background (a ceiling effect), making *Nc_med_diff* uninformative. ΔENC resolved this: *P/R* ΔENC medians were ~0.56–0.68 (near-zero relative to background), whereas background medians were ~11.46–11.76, yielding *DeltaENC_med_diff* ≈ −11 in all strains (all perm. $P$ = 1e−4). In clade B, effects were smaller yet consistent: *Fop_med_diff* ≈ −0.045 to −0.053 (all significant; perm. $P$ = 0.008–0.021), *Nc_med_diff* = 0 again by ceiling at 61, and *DeltaENC_med_diff* ≈ −2.35 to −3.32 (perm. $P$ = 2e−4 to 1.8e−3 ). Overall, *P/R* codon usage is closer to the composition-only expectations, whereas background genes deviate more, suggesting that reduced gBGC explains most of the signal, with at most modest relaxation of selection on synonymous codon usage.

#### (4) CAI after controlling for GC3/composition and CDS length

We observed that CAI values were lower for genes within the *P/R* locus than in background genes (e.g., median CAI = 0.231 vs. 0.397 in DSM108351; and 0.381 vs. 0.410 in CBS6039; **S4 Appendix**), raising the question of whether this difference reflects relaxed codon optimization or is merely a byproduct of nucleotide composition and gene length. CAI is a classic index of codon adaptation based on codon frequencies in highly expressed genes [8, 10], but it is confounded by background nucleotide composition; for instance, GC-rich third codon positions can inflate CAI even in the absence of selection. To address this, we tested whether the CAI deficit in *P/R* persists after accounting for composition and length using three complementary approaches.

We first applied an ordinary-least-squares (OLS) model to regress CAI on *inside_pr*, Δ(exon–intron GC; a gBGC-linked proxy for composition, defined as the difference between exonic and intronic GC within a gene), and log-transformed CDS length [CAI ~ inside_pr + Δ(exon-intron GC) + log(length_cds)]. We define "*inside_pr*" as a binary indicator equal to 1 for genes whose CDS lies entirely within the *P/R* locus (*pr_start* to *pr_end* on the same chromosome) and 0 for genes on the same chromosome outside that window; boundary-overlapping genes were excluded. The coefficient on *inside_pr* (reported as *ols_inside_beta*) estimates the CAI difference between *P/R* and background genes at fixed Δ and length**.** *ols_inside_beta* was < 0 in all species (significant in clade D and in 3 of 4 clade B species), indicating lower CAI in *P/R* after these controls.

Second, we trained the model CAI ~ Δ + log(length) on background genes only, then applied it to all genes to obtain residuals. Negative *resid_med_diff* values indicate that residuals for *P/R* genes were lower than background at equivalent Δ and length, with strong significance in clade D and weaker, species-dependent effects in clade B (see **S4 Appendix**).

Third, we performed a composition- and length-controlled comparison by matching each *P/R* gene to same-chromosome background genes within ±0.03 GC3 and ±10% CDS length and then tested paired CAI differences via sign-flip permutation (*CAI_match_median*/*CAI_match_p*; Fop analyzed analogously). Under this stricter design, median CAI differences were small and not significant (e.g., DSM108351 −0.013, $P \approx 1.0$; CBS6039 −0.002, $P = 1.0$), and matched Fop differences were likewise non-significant.

Together, these analyses indicate that most of the raw CAI deficit reflects underlying composition (GC3 and Δ[exon–intron GC]) and gene length, consistent with the ΔENC results, with at most a modest residual consistent with weak relaxation of selection on synonymous codon usage.

### Overall pattern and interpretation

Across both clades, the *P/R* locus showed lower GC, lower GC3, fewer optimal codons, and higher Nc, with consistently stronger signals in clade D. The neutral diagnostics (non-coding GC depletion and a shift toward AT3 in third codon positions) were pronounced in clade D and weaker in clade B, pointing to a larger reduction in gBGC in clade D. Selection-sensitive metrics agreed: Fop was lower inside *P/R*; Nc medians saturated at 61 (prompting reliance on ΔENC and CAI at matched GC3 and length); ΔENC was near composition-only expectations inside *P/R* (close to zero) but positive in background; and matched CAI differences were not significant. Taken together, the four complementary tests converge on reduced gBGC as the primary driver of the base-composition shift in the recombination-suppressed *P/R* locus, with at most a modest contribution from relaxed selection on synonymous codon usage.
