## Supplementary material for "Tracing the evolution and genomic dynamics of mating-type loci in *Cryptococcus* pathogens and closely related species": S2 Text

We considered the possibility that the fusion of the *P/R* and *HD* loci in *Cryptococcus* sp. 3 might be a consequence of a reduction in chromosome number, since this species possesses 13 chromosomes compared to the inferred ancestral 14-chromosome karyotype [1]. Under this scenario, linkage of the two *MAT* loci could have resulted from a chromosomal fusion between the ancestral *P/R*- and *HD*-bearing chromosomes, coincident with the loss of one centromere.

To evaluate this hypothesis, we first identified which centromere was lost in *Cryptococcus* sp. 3, strain CWM60451. Centromere positions were inferred computationally based on the presence of characteristic long terminal repeat (LTR) elements typically enriched at *Cryptococcus* centromeres, in combination with synteny comparisons and identification of the longest ORF-free intergenic regions [1-4]. These predictions were then cross-validated by synteny comparisons with closely related species. Using this approach, we traced the missing centromere to chr. 3 of *Cryptococcus* sp. 3, which corresponds to the *CEN4* region of *C. neoformans* H99 (**S9B Fig**). In *Cryptococcus* sp. 3, this region no longer exhibits the characteristic LTR-rich structure typical of functional centromeres, consistent with centromere inactivation and eventual loss. Comparative synteny analyses with *K. shandongensis* further suggest that this loss could have arisen through intercentromeric recombination, in which the fusion of two centromeres led to the inactivation of one and stabilization of the chromosome with a single functional centromere.

Importantly, the *MAT* locus in *Cryptococcus* sp. 3 is located on chr. 11 (**Fig 6A** and **S9 Fig**), which is unrelated to the chromosome where centromere loss occurred. Thus, the reduction from 14 to 13 chromosomes in this lineage cannot account for the colocation of *P/R* and *HD*. Instead, our results indicate that physical linkage of the two *MAT* loci in *Cryptococcus* sp. 3 must have arisen through a separate structural event, independent of karyotype reduction.
